# Distinct senescence mechanisms restrain progression of dysplastic nevi

**DOI:** 10.1101/2023.07.14.548818

**Authors:** Franziska K Lorbeer, Gabrielle Rieser, Aditya Goel, Meng Wang, Areum Oh, Iwei Yeh, Boris C Bastian, Dirk Hockemeyer

**Author notes:** Correspondence to: Boris Bastian, and Dirk Hockemeyer.

## Abstract

TERT promoter mutations (TPMs) are frequently found in different cancer types, including approximately 70% of sun-exposed skin melanomas. In melanoma, TPMs are among the earliest mutations and can be present during the transition from nevus to melanoma. However, the specific factors that contribute to the selection of TPMs in certain nevi subsets are not well understood. To investigate this, we analyzed a group of dysplastic nevi (DN) by sequencing genes commonly mutated in melanocytic neoplasms. We examined the relationship between the identified mutations, patient age, telomere length, histological features, and the expression of p16. Our findings reveal that TPMs are more prevalent in DN from older patients and are associated with shorter telomeres. Importantly, these TPMs were not found in nevi with BRAF V600E mutations. Conversely, DN with BRAF V600E mutations were observed in younger patients, had longer telomeres, and a higher proportion of p16-positive cells. This suggests that these nevi arrest growth independently of telomere shortening through a mechanism known as oncogene-induced senescence (OIS). These characteristics extend to melanoma sequencing data sets, where melanomas with BRAF V600E mutations were more likely to have *CDKN2A* inactivation, overriding OIS. In contrast, melanomas without BRAF V600E mutations showed a higher frequency of TPMs. Our data imply that TPMs are selected to bypass replicative senescence (RS) in cells that were not arrested by OIS. Overall, our results indicate that a subset of melanocytic neoplasms face constraints from RS, while others encounter OIS and RS. The order in which these barriers are overcome during progression to melanoma depends on the mutational context.

## Introduction

Dysplastic melanocytic nevi are initiated by mutations in proto-oncogenes activating the MAP-kinase pathway and are a risk factor and potential precursors for melanoma (1, 2). The hyperphysiological activation of MAP-kinase signaling triggers a wave of melanocyte cell divisions, which are then thought to be restrained from further proliferation and progression to melanoma by a process called oncogene-induced senescence (OIS) (3–6). The hyper-activation and fast proliferation leads to continuous DNA replication stress that can trigger cell cycle arrest and thereby incrementally depletes the pool of dividing cells and results in a stable lesion of growth-arrested cells (7, 8). It has been demonstrated that the two tumor suppressors p53 and pRB are major regulators of OIS and that the loss of cell cycle checkpoint genes such as *TP53* or *CDKN2A* can bypass this arrest (9–11). However, the strength with which individual mutations activate the MAPK pathways can differ. One of the most frequent and strongest mutations is BRAF V600E, compared to other mutations in BRAF or inactivating mutations such as loss of NF1, which are considered to be only weaker activating mutations of MAPK signaling (12, 13). These differences result in pathophysiological and molecular differences of nevi with BRAF V600E compared to those with other driver mutations. For example, nevi with BRAF V600E mutation are more frequently associated with distinct histological features such as more nested intraepidermal melanocytes, larger junctional nests, abrupt lateral circumscription, and larger cell size (14).

Recent sequencing efforts have revealed that there are intermediate melanocytic neoplasms with driver mutations in addition to the nevus initiating MAPK activating mutations, that fall short of melanoma. For example, mutations in the promoter of telomerase reverse transcriptase (TERT) have been observed in such melanocytic tumors, and thus can occur at the transition from nevus to melanoma, preceding the selection of mutations that deactivate cell cycle checkpoints (11, 15). Telomerase is responsible for maintaining chromosome ends and is required for the continuous proliferation of most human tissue stem cells and highly long-term proliferative cell types (16, 17). The so-called TERT promoter mutations (TPMs), which subvert the transcriptional silencing of *TERT* (18), are the most common non-coding mutation in cancer (19). In cells without telomerase expression, telomeres shorten continuously and due to the gradual erosion of telomeres, cells eventually enter replicative senescence. Thus, by triggering replicative senescence, telomere shortening functions as a strong tumor suppressor pathway, which leads to a permanent growth arrest (20). TPMs enable telomerase to be expressed and thereby provide a means for somatic cells to evade replicative senescence. TPMs become a selective advantage for cells whose telomeres are exhausted and have become limited in their proliferation (18).

As a subset of melanocytic nevi dysplastic nevi (DN) have defining characteristic clinical and histopathologic features. They are larger than most common acquired nevi and display a characteristic immune and stromal response (21, 22). While the initiating driver mutations and additional mutations of melanomas have been well cataloged by sequencing studies (12, 23, 24), the mutational landscape of DN, their telomere length and their senescence state have not been extensively characterized. Telomeres of telomerase-immortalized cancer cells are comparatively short, and initially TPMs are not sufficient to immortalize cells as additional cellular changes are required to fully stabilize the chromosome ends (18). One such additional change was recently described, where telomerase activity at telomeres is increased by mutations in the promoter of the *ACD* gene, which encodes the telomerase interactive protein TPP1 (25). These findings highlight that our understanding of the events required for the immortalization of cells and the physiological context in which TPMs undergo positive selection is still incomplete. By the analysis of small cohorts of melanocytic nevi it has been shown that TPMs can already be found in melanocytic nevi (26), however, most comprehensive studies of genomic aberrations of melanocytic and dysplastic nevi did not investigate TPMs as they analyzed only the coding region of the genome (27).

The early occurrence of TPMs in some melanocytic nevi poses a central challenge to our current understanding of the genetic makeup of DN and their progression to melanoma. If all nevi were effectively restrained by OIS, their cells would not undergo sufficient numbers of cell division to reach replicative senescence, and thus not require TPMs to support their immortalization. To resolve the role of telomeres and TPMs in early melanoma development, it is necessary to determine the genetic context in which telomeres become exhausted.

## Results

### TPMs are found in dysplastic nevi of older patients and in a mutually exclusive manner with BRAF V600E

To elucidate the genetic context of DN in which TPMs are selected, we sequenced exons and non-coding regions of genes frequently mutated in melanocytic neoplasms in a cohort of biopsy specimens of DN. The diagnosis was defined by histopathology, including only DNs deemed to have sufficient tumor cell content for analysis. Seventy-nine DN samples from patients of 9 to 87 years of age were successfully sequenced, using a custom targeted DNA sequencing approach of a panel of known melanoma-associated genes. Only cases with at least one pathogenic mutation in any of the regions of interest were included in subsequent analyses (Table S2, Fig. S1A, Supplemental Methods), as we could not rule out that cases without detectable mutations had insufficient tumor cellularity or sequencing coverage. In 74 cases (93.4%) a single mutation in the MAP-kinase pathway was detected (Fig. 1A). BRAF V600E was the most common alteration (n=50; 63.3%), and the average age of patients whose nevi carried this mutation was lower than those with other alterations (42.5 vs. 61.4 years, p<0.001) (Figs. 1B and S1B). When present, BRAF V600E was the only mutation identified. The melanocytic nevi without BRAF V600E mutation typically had alterative MAPK-pathway activating alterations such as in NRAS and KRAS as well as NF1 and KIT. In 31.0% (9 of 29) of them, more than one pathogenic mutation was identified, with TPMs being the most frequent additional mutation (8 of 9; 88.9%) (Fig. 1A, S1). In individual DNs, we also found rare pathogenic mutations in SF3B1, TET2, TP53 and CDKN2A (Fig.1A). Overall, the mutations found were in agreement with the mutation spectrum identified in previous studies of dysplastic nevi and melanoma (11, 12, 24).

**Fig. 1:**
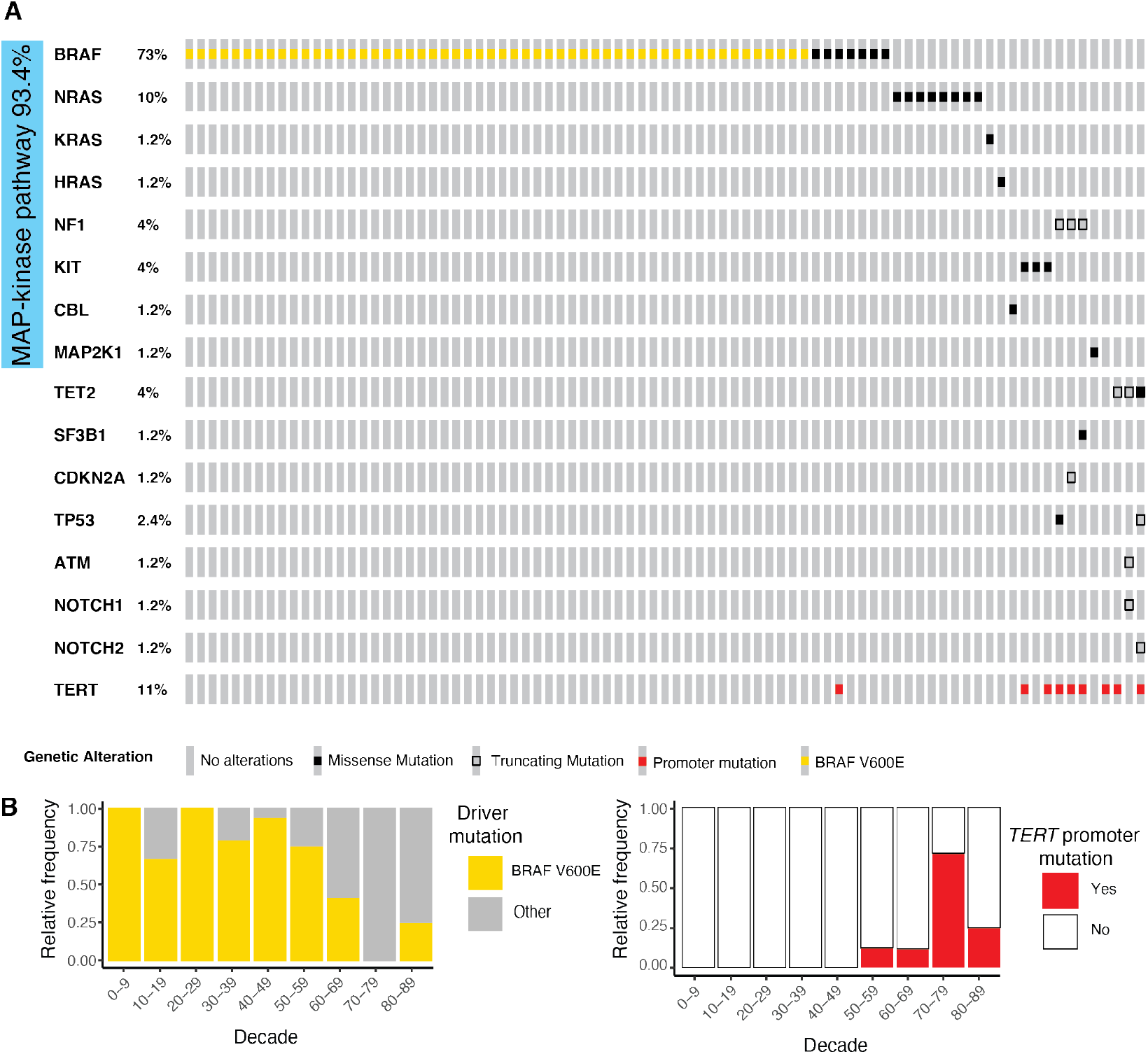
Somatic mutations in dysplastic nevi. BRAF V600E mutations are more common in younger patients where other mutations predominate in older patients, partially combined with *TERT* promoter mutations. (A) Tiling plot of pathogenic and likely pathogenic mutations. Blue bar indicates all mutations affecting the MAP-kinase pathway (B) Relative frequency of BRAF V600E (left) and *TERT* promoter mutations (right) by age decade.

### Telomeres remain long in DN with BRAF V600E mutations but shorten in those with other driver mutations

Since TPMs were exclusively detectable in samples from older patients (>54 years), we hypothesized that TPMs were selected because of age-related telomere-shortening. To address the question whether cells in DN with TPMs differed in the number of divisions they had undergone, we measured the relative telomere length of melanocytes in sections of the same DN samples. Comparisons of the telomere length differences in tissues from different individuals have to take into account the heterogeneity of telomere length within the human population. To overcome this challenge, we determined the telomere length of the melanocytes in the lesion relative to the surrounding tissue keratinocytes by developing a custom hybridization and image analysis pipeline (Figure 2A-C, Supplemental Methods). By normalizing the median intensity of telomeric signals from neoplastic melanocytes to that of the keratinocytes from the same section, we derived a “normalized telomere length” measurement, a procedure which largely eliminated variation in signal intensity between consecutive sections of cases and between individuals (Supplemental Figure 2C-H). We found that melanocytes in DN with TPMs had shorter telomeres than DN without TPMs (Figure 2D), supporting the notion that TPMs were selected once telomeres had eroded. On the other hand, we found that DN with BRAF V600E mutations had longer telomeres, whereas the DN with other driver mutations fell into two categories: 1) those with TPMs and very short telomeres, and 2) those without TPMs and with telomeres of intermediate length (p = 0.008, ANOVA) (Fig. 2E and Fig. S3). To compare the telomere lengths of DN to that of melanoma samples, we used melanoma samples known to carry TPMs from a prior study (18). We rank-ordered the telomere length distributions in the DN and compared them to those in the melanoma samples. We found that telomeres in DN with TPMs were or similar length or slightly longer than those in melanomas. In contrast, DN with BRAF V600E mutations had longer telomeres, indicating that these cells stopped dividing before their telomeres shortened (Figure 2D). In contrast, we concluded that DN with driver mutations other than BRAF V600E had undergone enough rounds of cell divisions for telomeres to have become eroded. In this context, TPMs were selected in a subgroup and could be placed in a proliferative trajectory closer to melanoma than samples without TPMs.

**Fig. 2:**
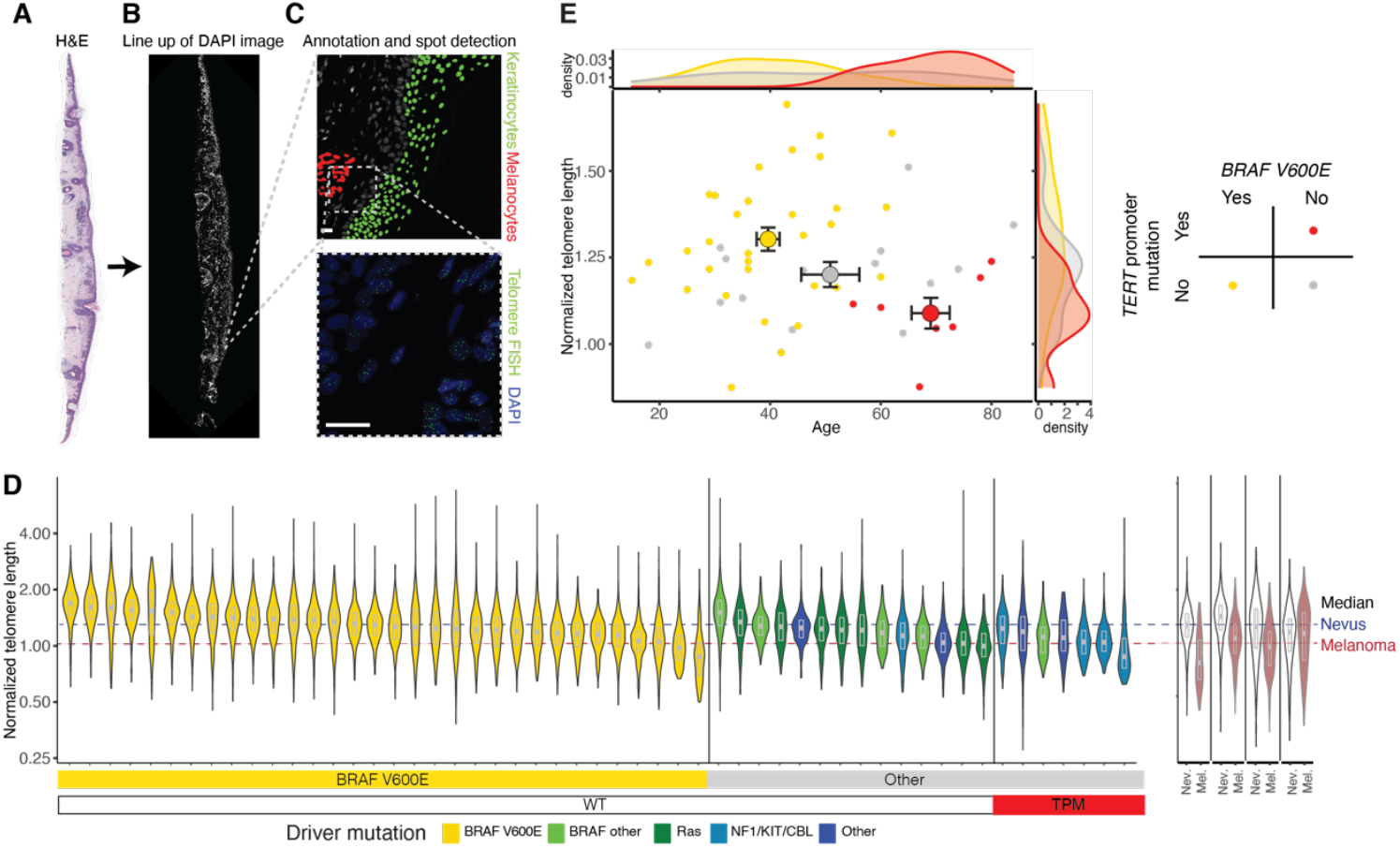
BRAFV600E mutant DN have longer telomeres than those without. Workflow for hybridization signal detection and identification of melanocytes and keratinocytes for quantitative telomere length analysis: (A) Scanning magnification of dysplastic nevus stained with hematoxylin and eosin. (B) Composite 4,6-diamino-2-phenylindole (DAPI) image of the same nevus. (C) Annotated DAPI image with melanocyte and keratinocyte population and their respective hybridization signals. (D) Normalized telomere length of the 53 successfully analyzed cases (n = 53) by BRAF V600E and TERT promoter mutation (TPM) status. The median telomere length of four matched adjacent nevus and melanoma cases shown on the right is indicated by the dashed lines for comparison. (E) Scatter plot and density distribution of age and median normalized telomere lengths of each nevus (small, filled circles) and group means ± standard error for each genotype (large, outlined circles).

### Dysplastic nevi with BRAF V600E mutations are histopathologically distinct and arrest with strong p16 induction

We then asked whether our cohort histopathologically recapitulates the differences, which have been observed in other cohorts dependent on the detected driver mutations. Independent and blinded scoring by two histopathologists revealed that the neoplastic melanocytes in DN with BRAF V600E mutations had a more nested distribution, with a larger proportion of cells in the dermis, whereas the other DN frequently had a lentiginous growth pattern and more subjacent solar elastosis (Fig. 3A, S4C). These results demonstrated that BRAF V600E DN presented a histopathologically distinct subgroup of DNs in our cohort, consistent with observations made in melanomas (28) and melanocytic nevi (14). Additionally, DN without BRAF V600E mutations were more frequently classified as neoplasms of uncertain behavior (ICD-10 D48.5) at the time of diagnosis (9 of 28; 34% vs 5 of 51; 9.8%, p= 0.028; Fisher’s Exact Test) than those with the mutation. Among DN with driver mutations other than BRAF V600E, DN with TPMs also had more frequent D48.5 codes compared to those without (6 of 9; 67% vs 8 of 70; 11.4%, p <0.001, Fisher’s Exact Test), indicating that they were histopathologically more atypical.

**Fig. 3:**
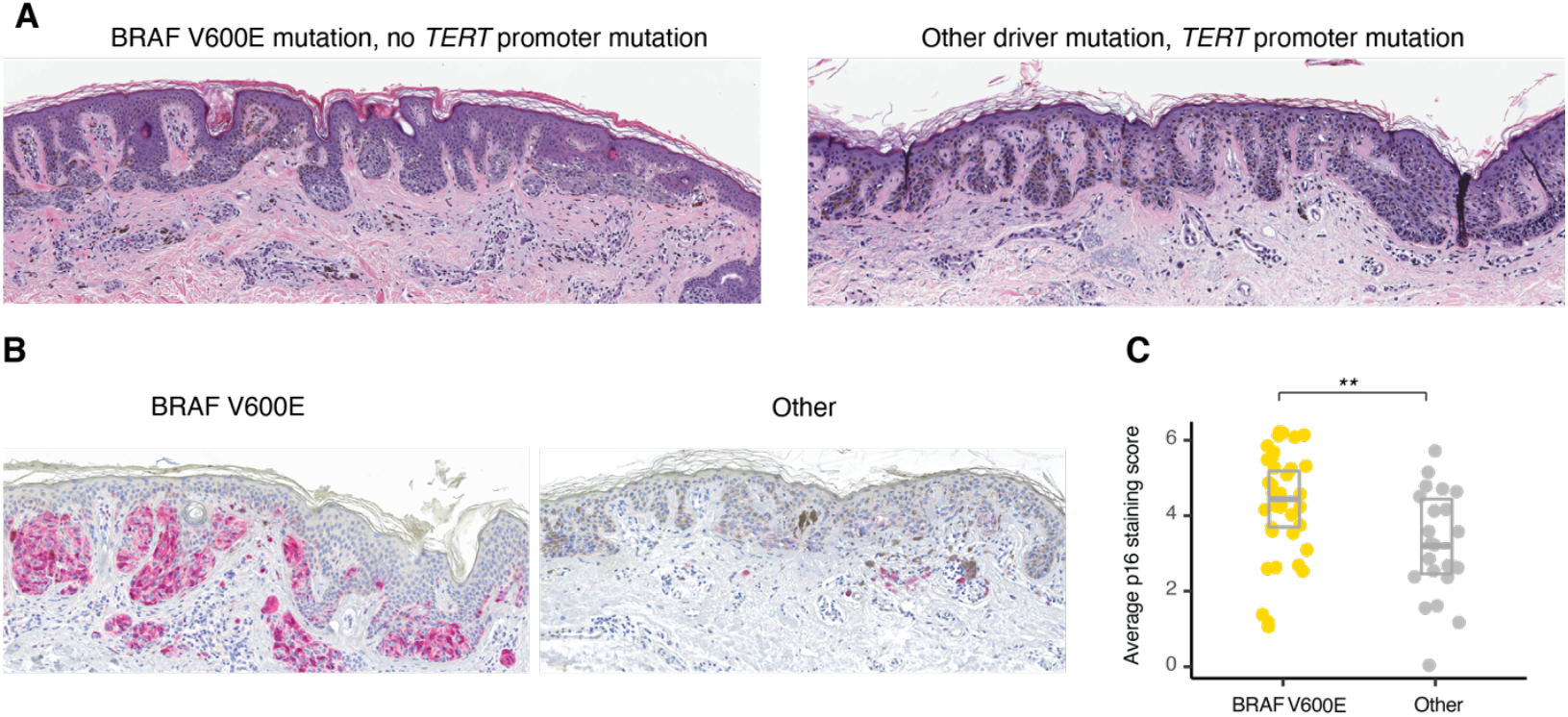
Dysplastic nevi with BRAF V600E mutation display higher levels of p16 expression by immunohistochemistry. (A) Photomicrographs of hematoxylin and eosin-stained dysplastic nevi with a BRAF V600E displaying a nested growth pattern (top) and without a BRAF V600E mutation and with a *TERT* promoter mutation displaying a lentiginous growth pattern with marked solar elastosis (bottom). (B) Immunohistochemistry for p16 (red) of a dysplastic nevus with (left panel) and without (right panel) BRAF V600E mutation. The right nevus also harbored a *TERT* promoter mutation. (C) Jitter plot with boxplot of average p16 immunoreactivity score of dysplastic nevi (n = 61, p = 0.005, Wilcoxon Rank Sum test).

Based on these findings, we tested the prediction that DN with driver mutations other than BRAF V600E also expressed less of the cyclin-dependent kinase inhibitor and tumor suppressor p16, because they are not arrested via OIS, and thus progressively shorten their telomeres. To test this, we evaluated the status of p16 in our DN cohort. We scored nuclear and cytoplasmic p16 immunostaining of DN sections. We found that DN with BRAF V600E expressed higher levels of p16 than those with other driver mutations (p=0.005, Kruskal-Wallis Rank Sum Test, Fig. 3, S4A). This difference was not dependent on the TPM status (Fig. S4B).

### Oncogenic driver mutations correlate with the mechanism of cellular immortalization in advanced melanoma

Our observations revealed that DN with BRAF V600E showed features distinct from DN with other driver mutations including longer telomere length and higher p16 expression. From this, we hypothesized that they would select subsequent mutations in a different sequential order, specifically melanomas arising from BRAF V600E mutant precursors would first have to override the oncogene-induced senescence checkpoint. To test the predictions of such an orthogonal senescence pathway model, we analyzed the available melanoma datasets for the co-occurrence of cell-cycle checkpoint inactivating mutations, which are required to overcome OIS (11, 12, 15, 24). Primary melanomas with BRAF V600E mutations indeed had more frequent inactivating mutations of the *CDKN2A* gene (45.2% vs 27.7%, p = 0.027, Fisher’s Exact test), consistent with early corruption of critical cell cycle checkpoint components triggered by OIS (Fig. 4A). Furthermore, primary melanomas with BRAF V600E mutations had fewer TPMs than those without (77.8% vs 93.8%, p = 0.024, Fisher’s exact test). This difference disappeared in metastases, indicating that TPMs arise after CDKN2A inactivation in these melanomas (Fig. 4B). Similar results were obtained when less frequent mutations of the G1/S checkpoint at the level of *CDK4* or *RB1* were also included in the analysis (Fig. S5). These findings indicated that in melanomas with BRAF V600E mutations, inactivation of the G1/S checkpoint typically arises before immortalizing mutations. In melanomas with other driver mutations, TPMs arose first whereas cell cycle checkpoint mutations emerged later.

**Fig. 4:**
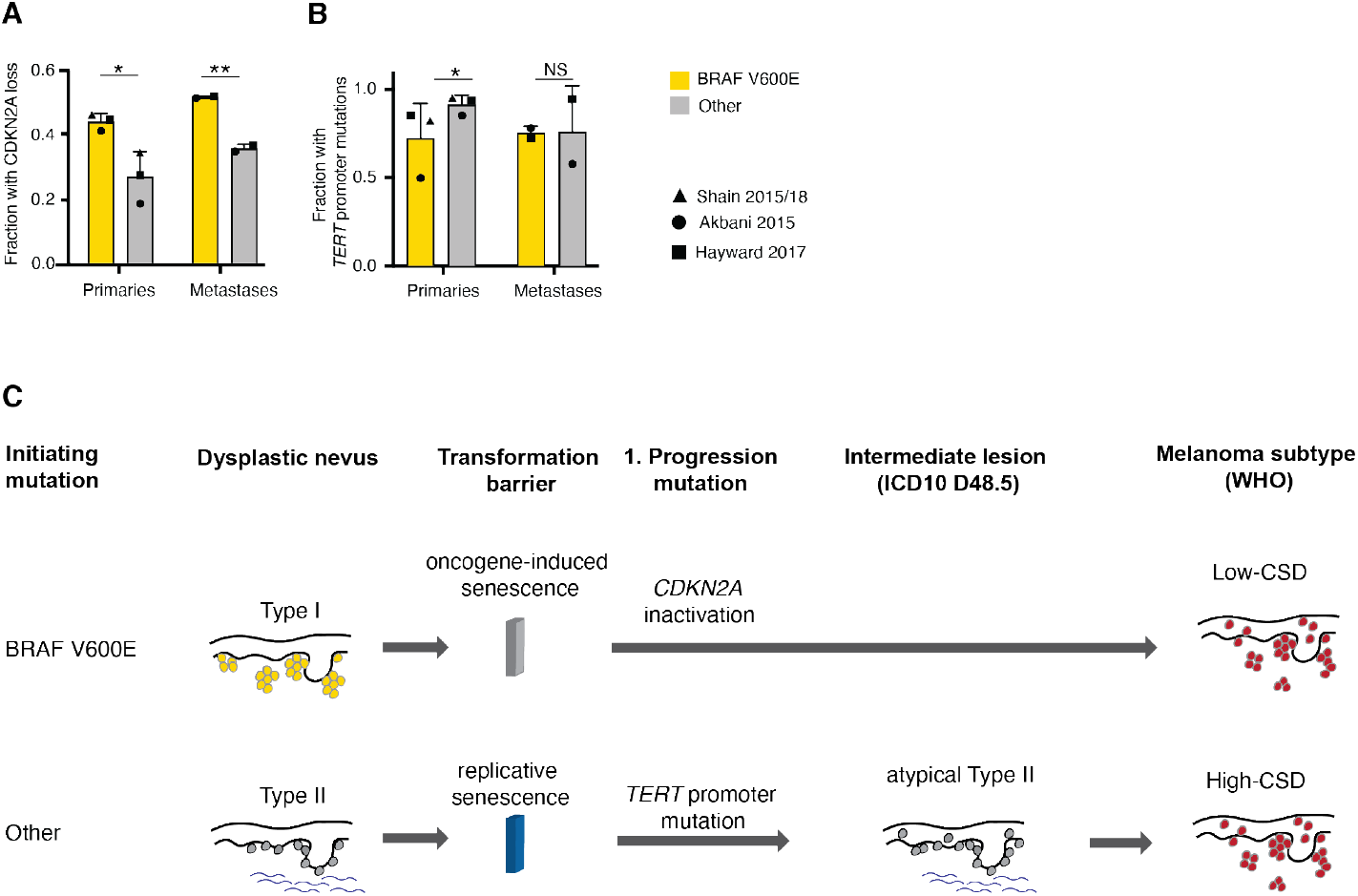
In primary melanomas with BRAF V600E mutations, biallelic inactivations of *CDKN2A* are more common and *TERT* promoter mutations are less common. Bar graph of the frequency of (A) biallelic *CDKN2A* inactivation (melanoma p = 0.027, metastasis p = 0.0076, Fisher’s exact test) and (B) *TERT* promoter mutations (melanoma p = 0.024, metastasis p = N.S., Fisher’s exact test) in primary metastases and melanoma metastases patients with and without BRAF V600E mutations. Bar graphs show mean and standard deviation of fraction of patients. Value of each individual study indicated by symbols. (C) Model of the evolutionary pathway of melanoma formation from the two proposed types of dysplastic nevi: The pathway to low-CSD melanomas (top) begins with a BRAF V600E mutation in a melanocyte, which clonally expands and forms a nevus. Cell proliferation is halted by the induction of oncogene-induced senescence, which arrests cells before they exhaust their telomeres. Transition to melanoma requires escape from oncogene-induced senescence, which, as the genomic data from primary melanomas indicate, frequently involves disruption of the *CDKN2A* locus (11, 15). The resulting loss of cyclin-dependent kinase inhibitors p16/INK4A p14/ARF and often p15/INK4B^40^ allows cells to resume proliferation, continue to shorten their telomeres, which ultimately selects for *TPMs* after formation of the primary melanoma. The pathway to high-CSD melanomas (bottom) begins with MAP-kinase pathway mutations other than BRAF V600E (11, 42). Nevi with these mutations have a more lentiginous growth pattern and more solar elastosis. In nevi with these mutations, oncogene-induced senescence is not triggered, and replicative lifespan constitutes the first transformation barrier. Some dysplastic nevi overcome this barrier by means of *TERT* promoter mutations and assume a more atypical histology. *CDKN2A* and equivalent mutations arise later during progression, *after TPMs* become prevalent.

## Discussion

By molecularly and genetically characterizing a prospectively collected set of DN, we found that DN, as defined by histopathologic features, are heterogeneous and can be divided into two main groups. Type I DN that are characterized by BRAF V600E mutations as the sole aberration and have an earlier age of onset. Their telomeres are long and their p16 expressions are similar to common acquired nevi, in keeping with the concept that they are predominantly kept in check by oncogene-induced senescence (5). Type II DN, are characterized by driver mutations other than BRAF V600E and are genetically more diverse. They have shortened telomeres, lower p16 expression levels, and frequently harbor TPMs, implicating replicative senescence as the first transformation barrier. These type II DN also have a more lentiginous growth pattern, whereas type I DN have a more nested growth pattern.

The categories of type I and II DN share key characteristics with the two main types of cutaneous melanomas defined in the current WHO Classification of Skin Tumors 6/29/23 4:12:00 PM(29). Specifically, type I DN share frequent BRAF V600E mutations, younger age, a more nested growth pattern, and lower degree of solar elastosis with melanomas on skin with low cumulative sun damage (CSD) (28). By contrast, type II DN are similar to high-CSD melanomas in that they lack BRAF V600E mutations, affect older individuals, have more solar elastosis, and a lentiginous growth pattern (30–32). Therefore, our data suggest that type I and II DN reside on two separate evolutionary trajectories, one leading to low- and the other to high-CSD melanoma, with disparate phasing of transformation barriers. A model illustrating the progression along the two pathways is shown in Figure 4C.

What could be the molecular underpinnings of the two different groups of DN? The finding that telomeres remain long in DN in younger patients, which mostly have BRAF V600E mutations suggests that telomeres shorten differently dependent on the driver mutation. It has been shown that different mutations in the MAP-kinase pathway activate the pathway to different degrees (13) and that high levels of activation can trigger OIS (33). Based on our observations, we conclude that the differences in telomere length in DN are the result of the differential activation of OIS and RS depending on the initiating driver mutation and represent two orthogonal senescence mechanisms limiting the progression of dysplastic nevi. However, an alternative explanation could be that the cell state of the melanocyte in which the driver mutation arises may also affect the fate of the nascent neoplasm and which tumor suppressor mechanisms become engaged, explaining the observed differences in telomere length and p16 accumulation (34, 35).

Genetic population studies of telomere length have correlated a longer constitutional telomere length to an increased cancer incidence rate (36). This observation is supported by rare familial melanoma mutations such as a TPM (37) or mutations in shelterin proteins POT1 and TIN2 that elongate telomeres (38–40). This effect of longer telomeres is likely because neoplasms, which are kept in check by replicative senescence can grow to larger sizes if the constitutional telomere length is longer. Thus the potential number of partially transformed cells arising from cell divisions increases and with that also the number of cells, that are at risk of acquiring mutations before telomere shortening limits their proliferation (41). We show that in addition to these consequences of constitutional telomere length, the telomere length changes following the acquisition of cancer initiating mutations impact the course of cancer development: A key novelty of our study is that we normalize for constitutional the telomere length using non-cancerous cells as a reference point. The analysis of relative telomere length in DN suggests that the observed length is the result of the differential activation of OIS and RS depending on the initiating driver mutation and that the two orthogonal senescence mechanisms limit the progression of dysplastic nevi. This measurement of relative telomere length supplements our characterization of the mutation status, the histopathology and the p16 status and enhances our understanding of the differences Type I and II DN as an early stage hyperplasia. Thus, our work provides a roadmap to determine if early lesions in other cancer types that frequently have a similar genetic progression - an initial MAPK oncogenic driver mutation, including BRAF V600E, followed by a TPM - can similarly be classified into two groups that differentially engage OIS and RS.

## Materials and Methods

### Patients

We prospectively collected DN based on histopathological features including a broad silhouette with lateral extension of the junctional component, eosinophilic or lamellar fibroplasia of the papillary dermis, or random cytologic atypia (42). Only lesions deemed to have sufficient tumor cell content for microdissection were included in the subsequent analyses. All lesions had received ICD10 codes of either *melanocytic nevus* (D22, n=65) or *neoplasm of uncertain behavior of skin* (D48.5, n=14) at diagnosis.

### Next-generation sequencing

DNA was extracted from microdissected sections and libraries were constructed for targeted next generation sequencing (NGS) of 80 genes implicated in melanocytic neoplasia (Supplementary Table 1), using previously reported methods of analysis (43). Only cases containing at least one pathogenic mutation in any of the targeted genes were included for further analysis, as we could not rule out that cases without any detectable mutations were false negatives due to insufficient tumor cell content (details in Supplemental Methods).

### Telomere length measurement

We performed quantitative fluorescence in situ hybridization (qFISH) for telomeric sequences as described (18). Hybridized tissue sections were scanned at 20x magnification by structured illumination microscopy on a Rebus Biosystems synthetic aperture optics custom microscope (44, 45). Images were spatially registered with hematoxylin and eosin stained sections to assist in identifying neoplastic melanocytes or epidermal keratinocytes (Fig. 2A-C) and analyzed without knowledge about the genotype of each case. Pixel intensity and area of individual hybridization signals were quantified using custom software (Rebus Biosystems). To mitigate the effects of inter-individual telomere length variation and varying hybridization efficiencies, we normalized the median signal intensity from neoplastic melanocytes to that of keratinocytes from the same section and derived a “normalized telomere length” measurement (Fig. S2, details in Supplemental Methods).

### Histomorphology assessment and Immunohistochemistry

Two dermatopathologists independently reviewed all cases and semi-quantitatively scored solar elastosis, the proportion of melanocytes in the dermis, and degree of nest formation of melanocytes, as previously described.(28) Solar elastosis was dichotomized into high (score ≥2) or low (score<2). The relative proportion of neoplastic melanocytes in the dermis, and their degree of nesting was assessed at scanning magnification and quantified as 1: <25%, 2: 25-50%, or 3: >50%. Interobserver agreement was assessed by a weighted kappa score and ranged from 0.48 to 0.86. Immunohistochemistry was performed on a Leica Bond instrument using a p16 antibody from Bio SB (Clone: 16P04, JC2) and the Leica Refine Red detection system. Immunoreactivity was scored blinded by two dermatopathologists separately for nuclear (0: no staining, 1: <25%, 2: 25-75%, and 3: > 75% positive nuclei) and cytoplasmic (0: for no, 1: weak, 2: moderate, and 3: strong) immunoreactivity. Both scores were added, and the two observers’ means used for statistical analysis.

### Re-analysis of existent melanoma datasets

We analyzed processed mutation data from of 512 cutaneous melanomas from the following publicly available datasets: Akbani et al. (TCGA),(46) Hayward et al.(47) and Shain et al.(15, 43) (details in Supplemental Methods).

## Statistical analysis

We used the Welch t-test for two-way comparisons of continuous variables (age, telomere length), ANOVA for comparisons of multiple groups followed by a posthoc Tukey’s HSD test, the Wilcoxon Rank Sum test and Kruskal-Wallis test for ordinal variables (nesting, dermal fraction, p16 score), and Fisher’s Exact test for dichotomous variables (solar elastosis, ICD-10 code, *CDKN2A* inactivation, checkpoint loss, TPM). Statistical tests and significance levels (* p<0.05, ** p < 0.01, *** p < 0.001) are indicated in the figure legends.

## Data availability

Sequencing data was deposited at in the Sequence Read Archive (SRA) under the accession number: PRJNA777884.

## Acknowledgments

The authors thank Drs., Daniel Pinkel and Alexander Stratigos, Michel DuPage and David Raulet for helpful comments on the manuscript. This work was supported by an NCI Outstanding Investigatory Award (1R35CA220481) to BCB. D.H. was supported by a Research Scholar Grants from the American Cancer Society (133396-RSG-19-029-01-DMC) by the Siebel Stem Cell Institute, the Chen Zuckerberg Biohub, and the Pew Charitable Trusts and the Alexander and Margaret Stewart Trust. F.K.L. received a Boehringer Ingelheim Fonds PhD Fellowship. The authors have no conflicts of interest to disclose.

## Figures and Tables

**Fig. S1:**
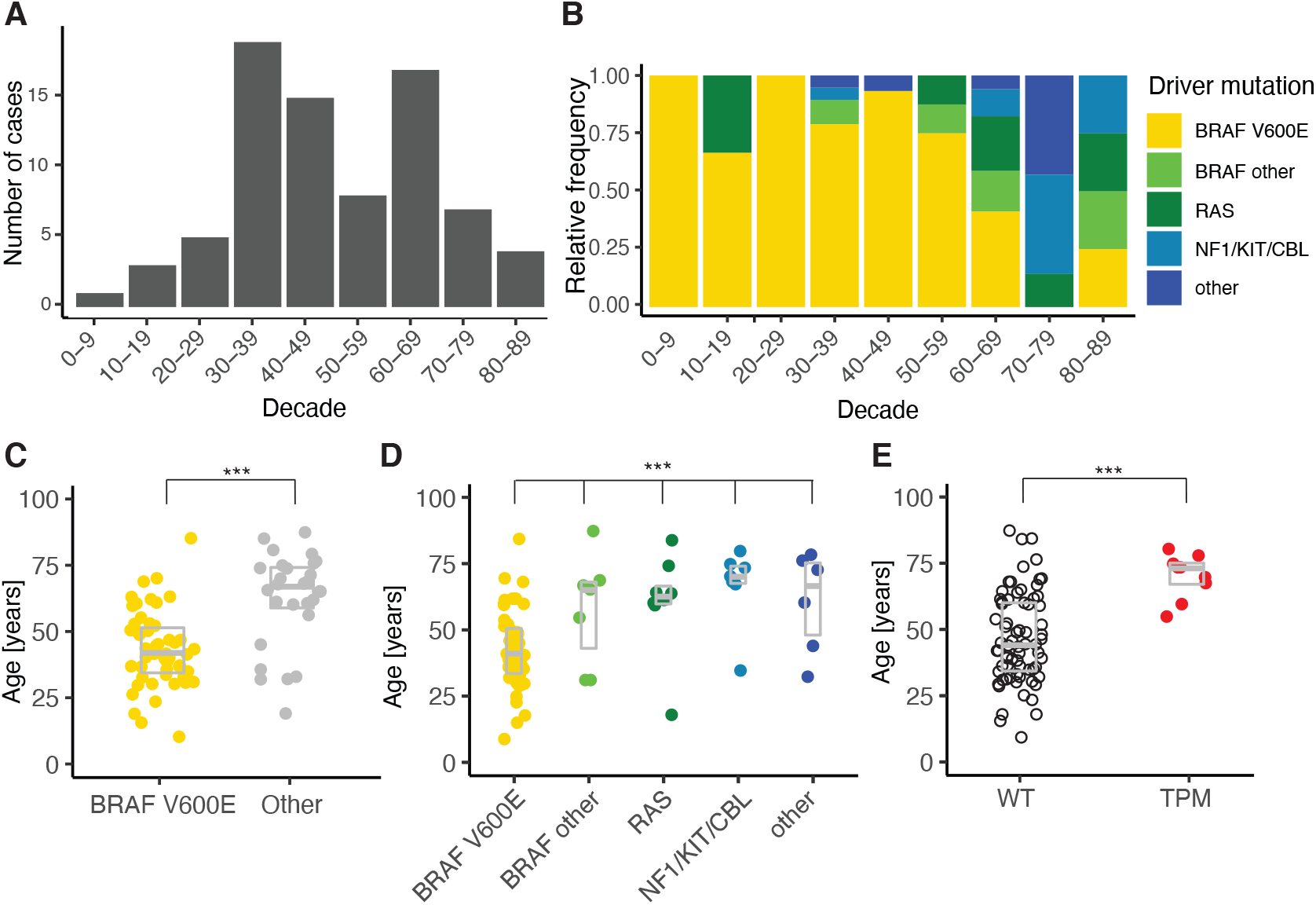
Age distribution of dysplastic nevi, their identified driver mutations and histopathological analysis. (A) Age distribution of the sequenced dysplastic nevi. (B) Relative frequency of driver mutations per age decade. (C) Patient age by BRAF V600E status (*** p<0.001, Welch’s t-test), (D) Individual subset of MAP-kinase pathway mutations (p<0.001, ANOVA), (E) *TERT* promoter mutation (TPM) status (p<0.001, Welch’s t-test).

**Fig. S2:**
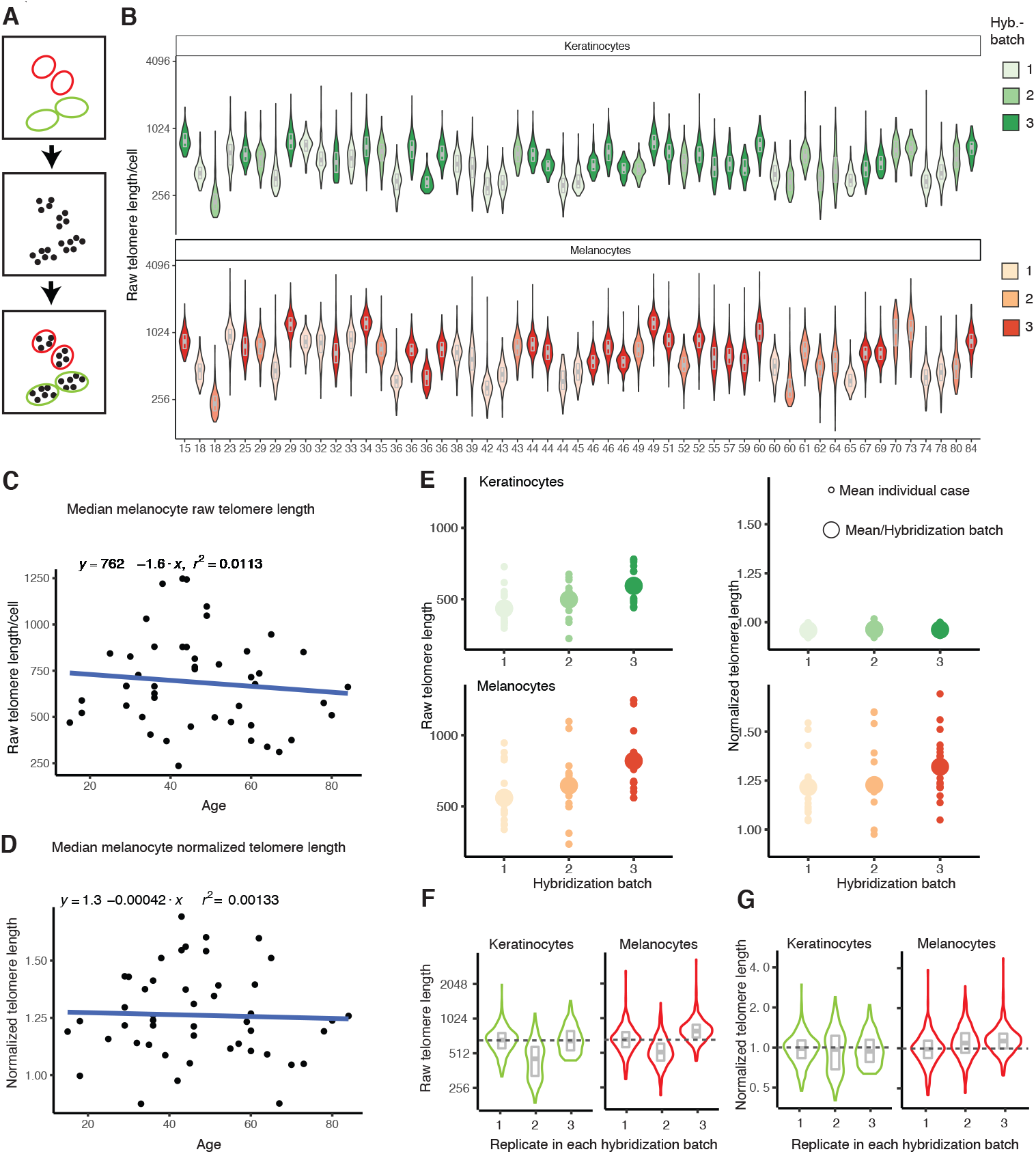
Normalization of melanocyte telomere length measurements to adjacent keratinocytes minimizes batch effects. (A) Schematic overview of the process of combining cell annotation information with hybridization signals to extract cell type-specific telomere length measurements. (B) Violin plots with inset boxplots showing median and quartiles of telomeric signals before normalization (red = melanocytes, green = keratinocytes, shades indicate staining batches). Scatter plot of raw (C) and normalized (D) median telomere length measurements of neoplastic melanocytes and patient age with linear regression line. (E) Median telomeric signal of melanocytes and keratinocytes in each nevus (small circle) and the average of all cases (big circle) for the three hybridization batches before (raw telomere length) and after normalization (relative telomere length). Violin plots with inset boxplots showing median and quartiles of raw (F) and normalized (G) telomeric signals of a dysplastic nevus stained in all three hybridization batches.

**Fig. S3:**
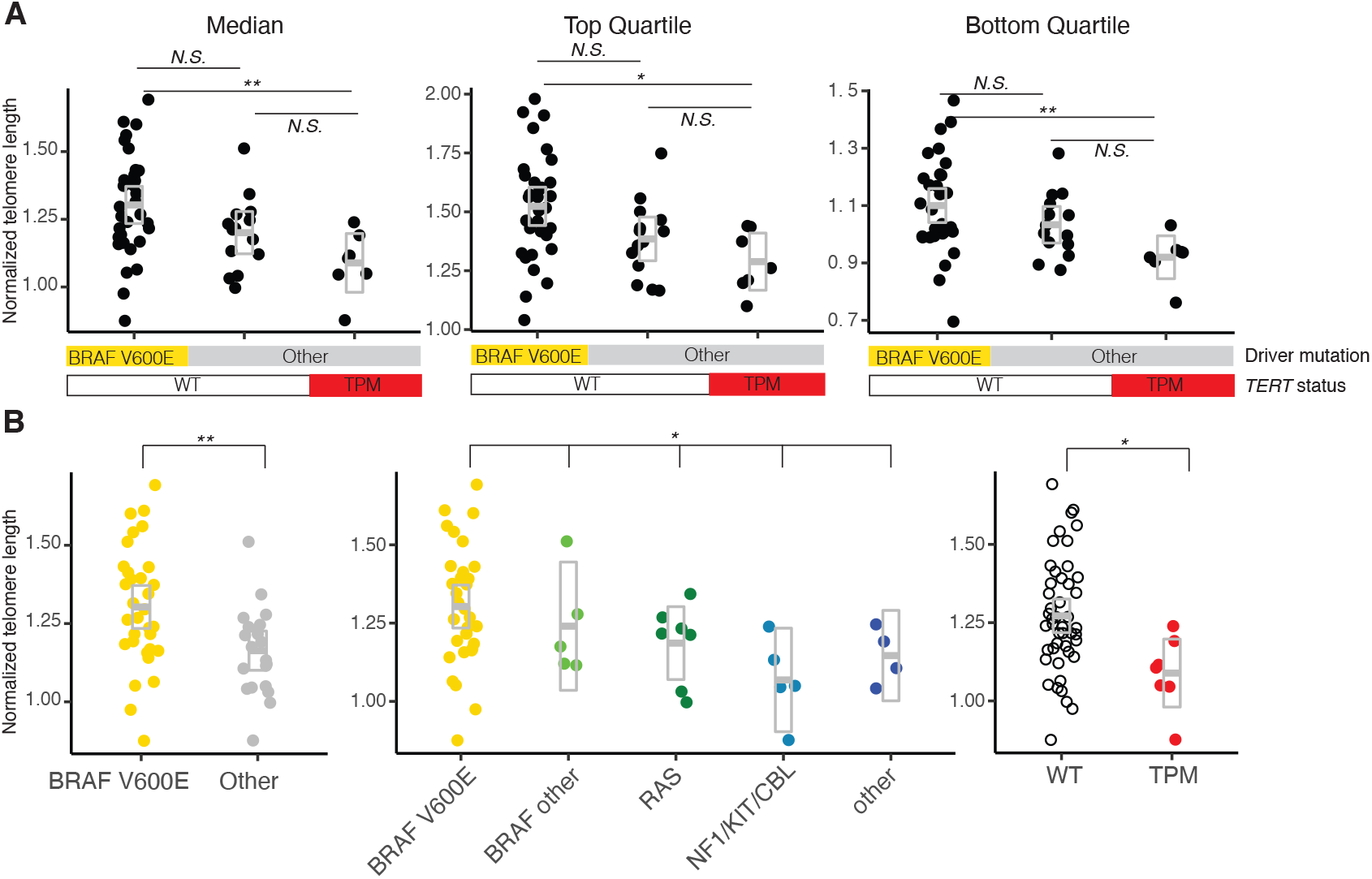
Telomere length in dysplastic nevi varies depending on MAP-kinase pathway driver and *TERT* promoter mutation status. (A) Jitter plots with inset boxplots of median, top and bottom quartile telomere lengths show that the cells of dysplastic nevi with BRAF V600E mutation tend to have longer telomeres over the entire distribution of telomere lengths within a given lesion (Median: p = 0.008, ANOVA, p = 0.011, Tukey multiple comparisons, Top quartile: p = 0.010, ANOVA, p = 0.019, Tukey HSD, Bottom quartile: p = 0.012, ANOVA, p = 0.011, Tukey HSD). (B) Jitter plots with inset boxplots of normalized telomere length compared by BRAF V600E mutation status (left panel, p = 0.006, Welch’s t-test), by groups of MAP-kinase driver mutations (middle panel, p = 0.039, ANOVA), and by *TERT* promoter mutation status (right panel, p = 0.013, Welch’s t-test).

**Fig. S4:**
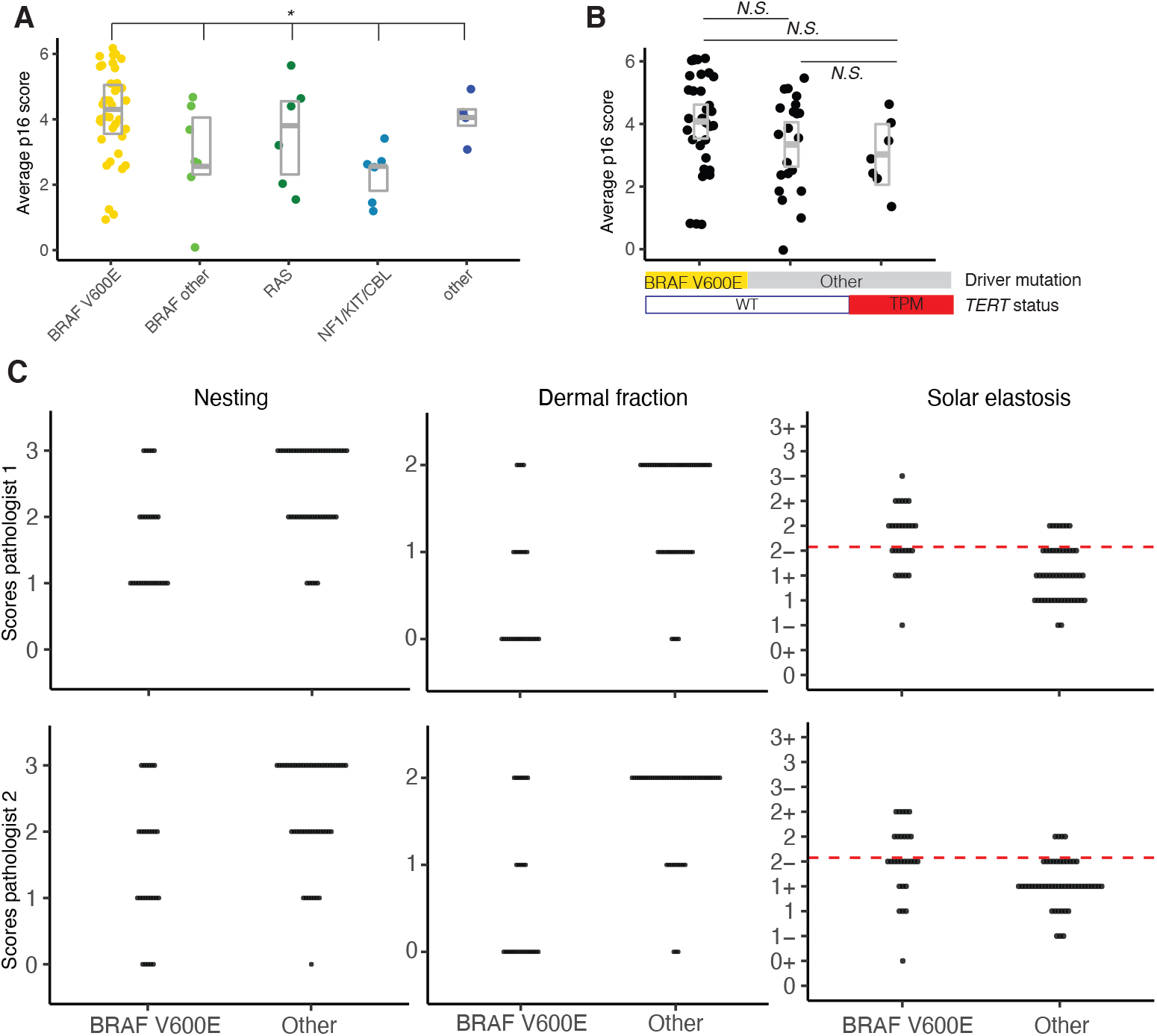
The p16 score of BRAF V600E mutant nevi is higher than of those with other driver mutations: (A) Jitter plot and boxplots of the average p16 score for all driver mutations (n = 61, p = 0.02, Kruskal-Wallis test) (B) Jitter plot and boxplots showing the average p16 score of all cases separated by BRAF V600E and *TERT* promoter mutation status(WT/TPM) status (n= 61, p, Kruskal-Wallis test). (C) Histopathological analysis of dysplastic nevi with BRAF V600E and other driver mutations scored by two pathologists: Nesting (Wilcoxon rank sum test, pathologist 1: p<0.001, pathologist 2: p<0.001, weighted kappa concordance coefficient: 0.51), Dermal fraction (Wilcoxon rank sum test, pathologist 1: p<0.001, pathologist 2: p = 0.005, weighted kappa concordance coefficient: 0.86), Solar elastosis (Fisher’s exact test High >=2, Low <2 (dashed red line), pathologist 1: p < 0.001, pathologist 2: p = 0.003, weighted kappa concordance coefficient: 0.30).

**Fig. S5:**
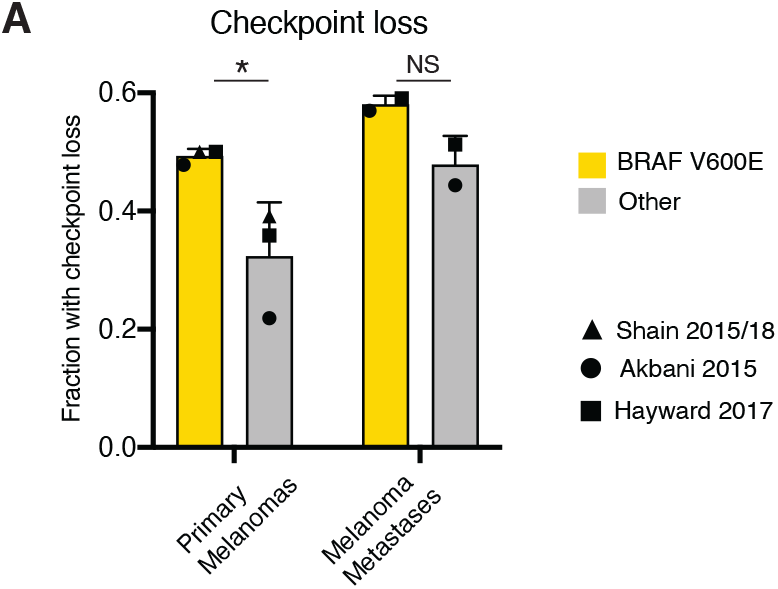
G1/S ceckpoint loss is more common in primary melanoma with driver mutations other than BRAF V600E: (A) Bar graphs showing the mean and standard deviation of the fraction of patients with G1/S checkpoint loss by BRAF V600E mutation status in primary melanomas and metastatic melanomas. Averages in individual studies are represented by different symbols (melanoma p = 0.042, metastasis N.S., Fisher’s exact test).

## Supplemental Methods

### Next-generation sequencing

After dissection and library construction libraries were pooled and captured with custom-designed bait libraries (xGen Lockdown probes, Integrated DNA Technologies, Coralville, Iowa) targeting the coding region of 80 genes known or suspected to be involved in the pathogenesis of cutaneous neoplasia and the promoter region of the *TERT* gene (Table S1). Sequencing was performed as paired-end, 100 base long reads on an Illumina HiSeq 4000 instrument (Illumina, San Diego, CA).

The reads were aligned to the human reference sequence UCSC build hg19 (NCBI build 37), using BWA-MEM 0.7.131.(Li and Durbin 2009) Variant calling was performed with Freebayes 0.9.20 and Unified Genotyper GATK.(Garrison and Marth 2012; McKenna et al. 2010) Variant annotation was performed with Annovar.(Wang et al. 2010) Copy number changes were called using CNVkit.(Talevich et al. 2016)

### Telomere length measurement

We performed quantitative fluorescence in situ hybridization (qFISH) using a Cy3-labeled probe for telomeric sequences as described previously.(Chiba et al. 2017) Hybridizations were performed in three randomized batches of samples. We scanned tissue sections counterstained with 4,6-diamino-2-phenylindole (DAPI) using structured illumination microscopy at 20x magnification using an *Esper* single cell multi-omics platform (Previously named StellarVision, Rebus Biosystems, Inc., Santa Clara, CA) based on Synthetic Aperture Optics, a structured illumination microscopy method.(Gustafsson 2000; Ryu et al. 2006)

In Synthetic Aperture Optics imaging, the sample was illuminated by a series of 12 high resolution light patterns formed by the interference of laser beams and the resulting series of low resolution images (“raw images”), which were then processed by *Esper* system’s on-board software (“*Esper Process*”) to generate a single high resolution image (“Synthetic Aperture Optics image”). This microscope allowed for an optical imaging resolution of a conventional 60x or 100x oil immersion microscopy at a low magnification and low numerical aperture air lens (20x/0.45NA), which offers more than an order of magnitude improvement in field of view, depth of field, and working distance compared to oil immersion lens. Based on this approach, we were able to image the entire tissue section with high resolution to measure the location and the intensity of telomere signals for each individual cell. Fluorescent spots representing telomere signals were detected by *Esper Process*, first by finding all local intensity peaks from the reconstructed Synthetic Aperture Optics image, followed by intensity thresholding to filter out the background. For each of the remaining spots, corresponding pixel location in the raw images was identified and the average value and the modulation strength of the series of 12 intensities of the pixel were calculated, based on which false positive spots were further eliminated.

Individual images were stitched together and spatially registered with a digital image of a hematoxylin and eosin (HE) stained section to assist in identifying regions of interest containing neoplastic melanocytes or epidermal keratinocytes (Fig. 2A-C). Individual nuclei were segmented based on the DAPI image using the implementation of Huang’s fuzzy thresholding in Fiji to turn DAPI images into binary images and the segmented nuclei were subsequently color coded as melanocytes or keratinocytes using morphologic and positional information from the HE images without knowledge about the genotype of each case (Fig. 2B, Fig. S2A). The minimum number of melanocytes quantified was 284 with an average number of 1265.4 melanocytes evaluated per case.

Based on their position, the telomeric signals were then assigned to individual keratinocytes and melanocytes and the median telomeric signal per cell was calculated (Fig. S2A-B). Telomere length varies among individuals, partially dependent on age and constitutional telomere length.(Allsopp et al. 1992; Rufer et al. 1999) Additionally, the intensity of hybridization signals can vary depending on tissue fixation, hybridization conditions, and batch effects (Fig. S2B,D). To mitigate both effects on the telomere length measurement, we normalized the median signal intensity from neoplastic melanocytes to that of the keratinocytes from the same section and derived a “normalized telomere length” measurement. This procedure largely eliminated variation in hybridization intensity between consecutive sections of cases (Fig. S2C-H) and eliminated inter-individual variation of telomere length. Data analysis was performed in R and plots generated with ggplot2.

### Re-analysis of existent melanoma datasets

We analyzed processed mutation data from of 512 cutaneous melanomas from the following publicly available datasets: Akbani et al. (TCGA)(Akbani et al. 2015), Hayward et al.(Hayward et al. 2017) and Shain et al.(Shain et al. 2015, 2018). Copy number data were obtained from the supplementary material(Shain et al. 2015; Hayward et al. 2017; Shain et al. 2018), from the Genomic Data Commons data portal of the National Cancer Institute(Akbani et al. 2015), or recalculated from the BAM alignment files using FACETS(Krauthammer et al. 2015; Shen and Seshan 2016). Samples that were from cell lines, could not be unequivocally classified as primary or metastasis, or had insufficient tumor cell content to derive copy number variation profiles were excluded, leaving 482 cases for further analyses.

Samples that had either inactivation of *CDKN2A* or *RB1* or had activation of CDK4 were considered to have an inactivated G1/S cell cycle checkpoint. *CDKN2A* and *RB1* were considered inactivated, if their loci displayed a deep deletion, or their coding sequences had a pathogenic mutation accompanied by either a shallow deletion, loss of heterozygosity, or an additional pathogenic mutation. Four samples with somatic mutations of unknown significance of *CDKN2A* (F90L, L65LL insertion, M52R and A57S) were excluded from the analyses. Activation of CDK4 was defined by the presence of a known activating mutation or amplification of the *CDK4* locus. *TERT* promoter mutation status could be derived from 263 samples that had sufficient sequencing coverage of the *TERT* promoter.(Shain et al. 2015)

